# PvC3H29 interacts with and inhibits DNA binding of PvNAPs to finetune leaf senescence in switchgrass (*Panicum virgatum*)

**DOI:** 10.1101/2022.04.14.488389

**Authors:** Zheni Xie, Guohui Yu, Shanshan Lei, Hui Wang, Bin Xu

## Abstract

Finetuning the progression of leaf senescence is important for plant’s fitness in nature, while ‘staygreen’ with delayed leaf senescence has been considered as a valuable agronomic trait in crop genetic improvement. In this study, a switchgrass CCCH-type Zinc finger gene, *PvC3H29*, was characterized as a suppressor of leaf senescence that over-expressing or suppressing the gene led to delayed or accelerated leaf senescence, respectively. Transcriptomic analysis marked that chlorophyll catabolic pathway genes were involved in the PvC3H29-regulated leaf senescence. PvC3H29 was a nucleus-localized protein with no transcriptional activity. By Y2H screening, we identified its interacting proteins, including a pair of paralogous transcription factors, PvNAP1&2. Over-expressing the *PvNAPs* led to precocious leaf senescence at least partially by directly targeting and transactivating chlorophyll catabolic genes to promote chlorophyll degradation. PvC3H29, through protein-protein interaction, repressed the DNA-binding efficiency of PvNAPs and alleviated its transactivating effect on downstream genes, thereby functioned as a ‘brake’ in the progression of leaf senescence. Moreover, over-expressing *PvC3H29* resulted in up to 47% higher biomass yield and improved biomass feedstock quality, reiterating the importance of leaf senescence regulation in the genetic improvement of switchgrass and other feedstock crops.

**One-sentence summary:** PvC3H29 interacts with transcription factors, PvNAP1&2, to inhibit their transactivation on chlorophyll catabolism and leaf senescence in switchgrass.

## Introduction

The progression of leaf senescence is finely modulated in plants. On one hand, leaf senescence is necessary for plants to recycle nutrient to new sink organs; on the other hand, precocious leaf senescence shortens the photosynthetic period and limits grain or whole biomass yield. For cereal, forage, and bioenergy crops, delayed leaf senescence or functional ‘staygreen’ is often considered as an important agronomic trait (Yang and Udvardi, 2018; Shin *et al*., 2020).

One key feature of the ‘staygreen’ trait was the retention of green appearance in aging leaves with delayed deconstruction of the photosynthetic apparatus (Thomas and Howarth, 2000). Chl degradation is a key step in leaf senescence that is catalyzed in the PAO (Pheophorbide a Oxygenase)/phyllobilin pathway involving a set of Chl catabolic enzymes (CCEs), including NYC1 (NON-YELLOW COLORING 1, the Chl *b* reductase), NOL (NYC1-like), HCAR (Hydroxymethyl Chl a Reductase), SGR/NYE1 (STAYGREEN or Non-Yellowing Protein 1, the Chl a Mg-dechelatase), PPH (PHEOPHYTINASE), PAO, and RCCR1 (Red Chl Catabolite Reductase) (reviewed by Kuai *et al*., 2018). Expression of these *CCGs* are tightly controlled during leaf aging to avoid drastic Chl degradation under normal growth condition. On the other hand, finetuning expression levels of *CCG*s showed promises in crop genetic improvement (Zhou *et al*., 2011; Shin *et al*., 2020; Yu *et al*., 2021).

Plants employed a set of transcription factors to transactivate or repress the expression of these *CCG*s. For example, NAP (NAC-LIKE, ACTIVATED BY AP3/PI, NAC029) was one transcription factor targeting on *CCG*s and other senescence-promoting genes to accelerate leaf senescence in different plant species, such as Arabidopsis (*Arabidopsis thaliana*), cotton (*Gossypium hirsutum*), rice (*Oryza sativa*), and bamboo (*Bambusa emeiensis*) (Chen *et al*., 2011; Fan *et al*., 2015; Yang *et al*., 2014; Liang *et al*., 2014; Sakuraba *et al*., 2014). In rice, expression of *OsNAP* was tightly linked with the onset of leaf senescence in an age-dependent manner and was induced specifically by abscisic acid (ABA); and OsNAP directly targeted genes related to Chl degradation (i.e., *SGR, NYC1, NYC3/PPH*, and *RCCR1*) and several other senescence associated genes (*SAGs*) (Liang *et al*., 2014). In Arabidopsis, Yang *et al*. (2014) found that NAP could directly target the ABA-biosynthesis gene, *ABSCISIC ALDEHYDE OXIDASE3 (AAO3),* to increase ABA content. One recent study showed that NAP also acted as an indispensable regulator in GA (gibberellic acid)-mediated leaf senescence that two DELLA proteins, namely GA-INSENSITIVE (GAI) and REPRESSOR OF ga1-3 (RGA), interacted with NAP, and the interaction subsequently impaired the transcriptional activities of NAP to induce the expression *SAG113* and *AAO3* (Lei *et al*., 2020). Furthermore, reducing the expression of *NAP* showed promises in crop genetic improvement. For examples, down-regulation of *GhNAP* significantly delayed leaf senescence and improved 15% of lint yield in cotton (Fan *et al*., 2015), and reduced *OsNAP* expression delayed leaf senescence and extended grain-filling period resulting in a 6.3-10.3% increase in rice grain yield (Liang *et al*., 2014). In short, the current working model pinpointed that NAP perceived ABA and GA signals and further accelerated leaf senescence by activating ABA-biosynthetic and Chl catabolic genes in a trifurcate feedback loop. Till now, it is unclear whether there is a ‘brake’ regulator targeting on NAP to delay the rate of leaf senescence.

CCCHs is a unique subfamily of zinc finger proteins, and several CCCHs were known as negative regulators in leaf senescence, such as *OsDOS* (Kong *et al*., 2006) and *OsTZF1* (Jan *et al*., 2013) in rice *(Oryza sativa), GhTZF1* in cotton (*Gossypium hirsutum*) (Zhou *et al*., 2014), and *PvC3H69* in switchgrass (*Panicum virgatum*) (Xie *et al*., 2021). Over-expressing of these *CCCHs* delayed leaf senescence with altered expression of genes involved in reactive oxygen species (ROS) homeostasis and ABA/JA signaling pathways (Kong *et al*., 2006; Jan *et al*., 2013; Xie *et al*., 2021). Yet, how exactly these CCCHs functioned in the regulation of leaf senescence is unclear. Furthermore, zinc finger proteins make tandem contacts with their target molecules, such as DNA, RNA, proteins, and lipids through their Zinc finger (Znf) motifs. It remains unknown whether there was a link between these CCCH proteins and senescence-promoting transcription factors (e.g. NAP) to finetune the senescence process.

Switchgrass is a tall perennial C4 grass species dedicated for bioenergy and forage feedstock production (Anderson *et al*., 1988; McLaughlin and Kszos, 2005). ‘Stay-green’ is a highly desirable trait for the tall grass. In this study, we identified one CCCH-type protein, PvC3H29, as a negative regulator in leaf senescence. PvC3H29 *per se* had no transcriptional activity but physically interacts with a pair of paralogous NAPs (PvNAP1&2). The PvNAPs promoted leaf senescence at least partially by directly targeting and transactivating *CCGs*. Through the interaction, PvC3H29 effectively attenuated the DNA binding efficiencies of the PvNAPs to delay the progression of leaf senescence. Furthermore, over-expressing *PvC3H29* resulted in up to 47% higher biomass yield in switchgrass. Thus, results of this study revealed a new regulatory module in leaf senescence and demonstrated the effectiveness of leaf senescence regulation in biomass improvement in switchgrass.

## Results

### PvC3H29 is a negative regulator in leaf senescence

To investigate the function of PvC3H29, we generated both over-expression and RNA-interference switchgrass transgenic lines (abbreviated as OE29 and RNAi lines, respectively) using the *Agrobacterium*-mediated transformation. The OE29 and RNAi lines had 6-9 times higher or 50-60% lower expression levels of *PvC3H29* than the wild type (WT) plants, respectively (Supplementary Fig. S1).

Phenotypically, the OE29 lines displayed a greener appearance than the WT (Fig 1a; Supplementary Fig. S2). At the ‘R3’ flowering stage of switchgrass (Hardin *et al*., 2013), leaves of OE29 lines showed greener appearance, higher Chl contents, and higher photochemical efficiencies (Fv/Fm) than those of WT (Supplementary Fig. S2). Taking detached leaves for dark-induced leaf senescence, it showed that Chl contents and Fv/Fm of WT leaves declined to less than 10%, while those of OE29 lines still retained over 70% after 15 days of dark treatment (DAD) (Fig.1A-C).

**Fig. 1.**
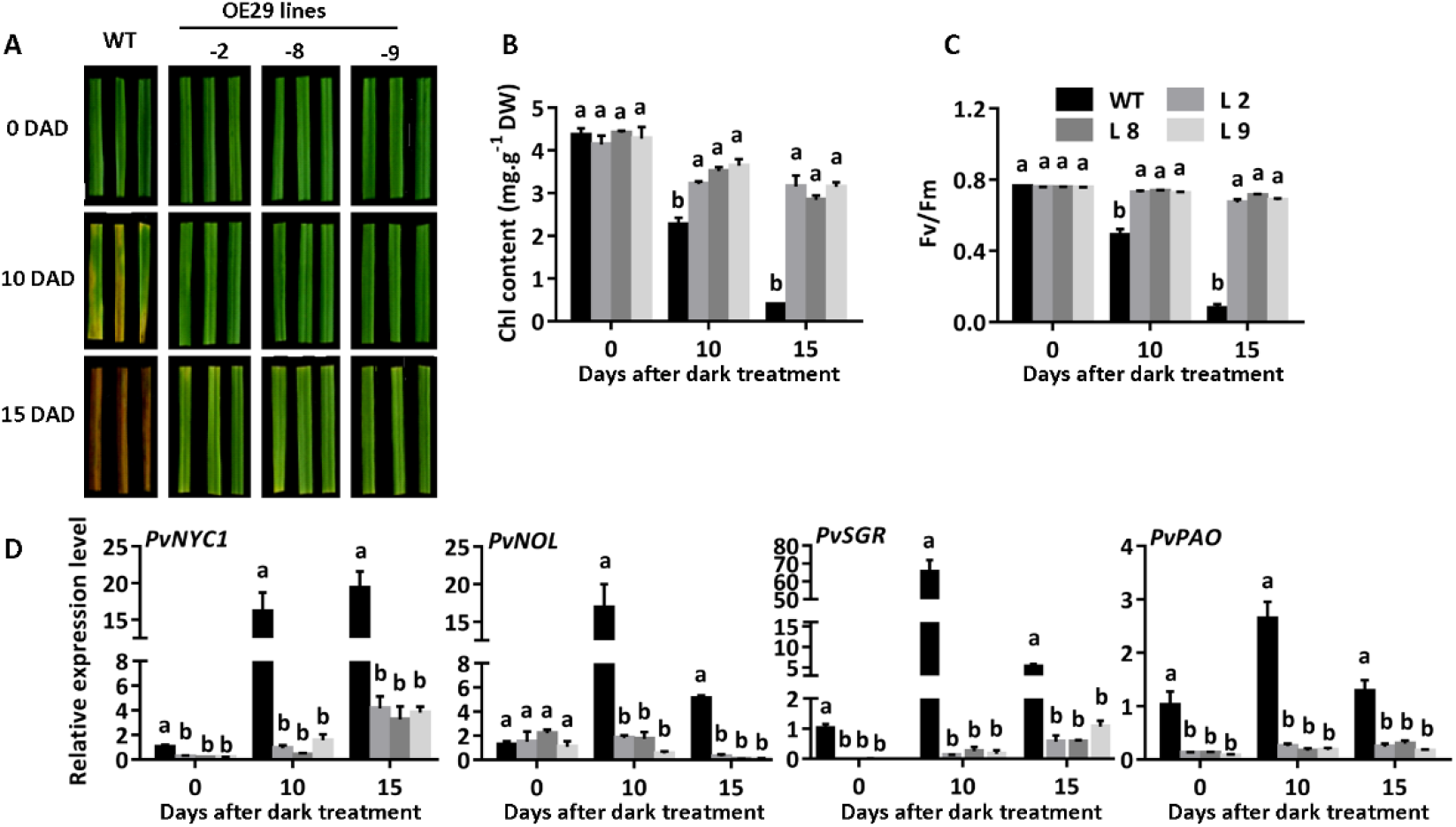
Over-expressing *PvC3H29* delayed leaf senescence in switchgrass. A, Phenotype of WT and OE29 transgenic lines. Detached green leaves at the same developmental stage were placed in dark for 15 DAD (<u>d</u>ays <u>a</u>fter <u>d</u>ark treatment). **B-D,** Chl contents. Fv/Fm values, and relative expression of four Chi catabolic genes (*PvNYC1, PvNOL, PvSGR*, and *PvPAO*) after 10 and 15 DAD in WT and OE29 lines. Letters above bars indicate significant difference at *P* < 0.05.

Consistently, relative expression levels of *CCGs* (e.g., *PvNYC1, PvNOL*, *PvSGR*, and *PvPAO*) in OE29 lines were significantly lower than those in WT (Fig.1D).

In contrast to OE29, the RNAi lines manifested precocious leaf senescence. Compared with WT, RNAi lines showed earlier and severer leaf chlorosis with significantly lower Chl contents, lower Fv/Fm values, but higher expression levels of *PvNYC1, PvNOL, PvSGR,* and *PvPAO* at 6 and 10 DAD (Fig. 2). Together, these results showed that PvC3H29 was a negative regulator in leaf senescence.

**Fig. 2.**
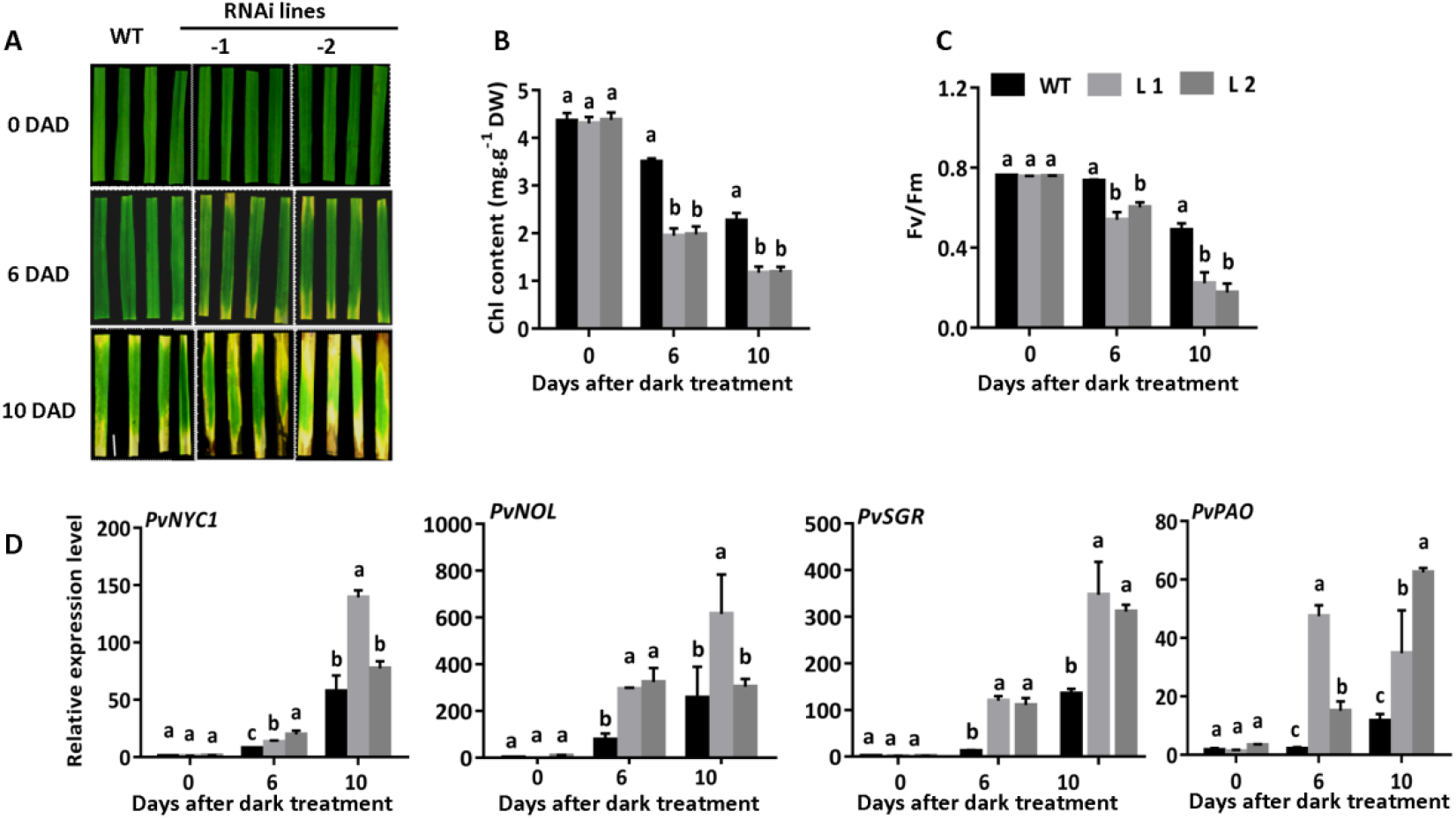
Suppressing *PvC3H29* accelerated leaf senescence in switchgrass. A, Phenotype of WT and *PvC3H29*-RNAi lines. Detached green leaves at the same developmental stage were placed in dark for 10 DAD (<u>d</u>ays <u>a</u>fter <u>d</u>ark treatment). B-D. Chl contents, Fv/Fm values, and relative expression of four Chl catabolic genes (*PvNYCl, PvNOL, PvSGR*, and *PvPAO*) after 6 and 10 DAD in WT and RNAi lines. Letters above bars indicate significant difference at *P* < 0.05.

### Transcriptomic comparisons among WT, OE29, and RNAi transgenic lines

Transcriptomic comparisons among WT, OE29, and RNAi transgenic lines were carried out using detached leaves at 7 DAD. As shown in supplementary figure S3, there were a total of 14,027 and 15,285 differentially expressed genes (DEGs) in ‘WT-7-vs-OE29-7’ and ‘WT-7-vs-RNAi-7’. For both ‘‘WT-7-vs-OE29-7’ and ‘WT-7-vs-RNAi-7’, Gene Ontology (GO) analysis of the DEGs showed highly similar results that the top two enriched ones were ‘catalytic activity’ and ‘binding’ in the ‘molecular function’ category, ‘metabolic process’ and ‘cellular process’ in the ‘biological process’ category, and ‘cell’, ‘cell part’ and ‘organelle’ in the ‘cellular component’ category (Supplementary Fig. S4). Kyoto Encyclopedia of Genes and Genomes (KEGG) pathway analysis showed that the top two enriched pathways were ‘metabolic pathway’ and ‘biosynthesis of secondary metabolites’ for both ‘WT-7-vs-OE29-7’ and ‘WT-7-vs-RNAi-7’ (Supplementary Fig. S5). Furthermore, we found that in the comparison of ‘WT-7-vs-OE29-7’, five out of seven key *CCG*s showed lower expression levels in OE29-7; while in ‘WT-7-vs-RNAi-7’; six out of seven *CCG*s (except *NYC1)* showed higher expression levels in ‘RNAi-7’ (Supplementary Fig. S6).

### PvC3H29 is a nuclear-localized protein with no transcriptional activity

Subcellularly, PvC3H29 was a nuclear-localized protein that the fluorescent signal of PvC3H29-GFP was merged with the DAPI-stained nuclear signal in ryegrass protoplasts (Fig. 3A). Using the yeast auto-transactivation assay, we found that neither PvC3H29 nor the negative control (GUS) infusion with the GAL4 DNA-binding domain (GAL4-DB) activated the reporter gene, while the positive control PvC3H72 did (Xie *et al*., 2019) (Fig. 3B). Furthermore, we used the GAL4-DB and its binding sites (GAL4[4×]-D1-3[4×]-GUS)-based *in planta* transient expression system to detect whether PvC3H29 was a transcriptional repressor. As shown in figure 3C, the PvC3H29 had no transcriptional effect on the reporter gene (Fig. 3C). Furthermore, the expression of *PvC3H29* was 3-5 times higher in the 2^nd^-5^th^ leaves from the top than in the 1^st^ leaf (the flag leaf) at the R3 stage of switchgrass (Hardin *et al*., 2013). Together, these results showed the expression of *PvC3H29* was related to leaf ages, and PvC3H29 was a nucleus localized protein with no transcriptional activity.

**Fig. 3.**
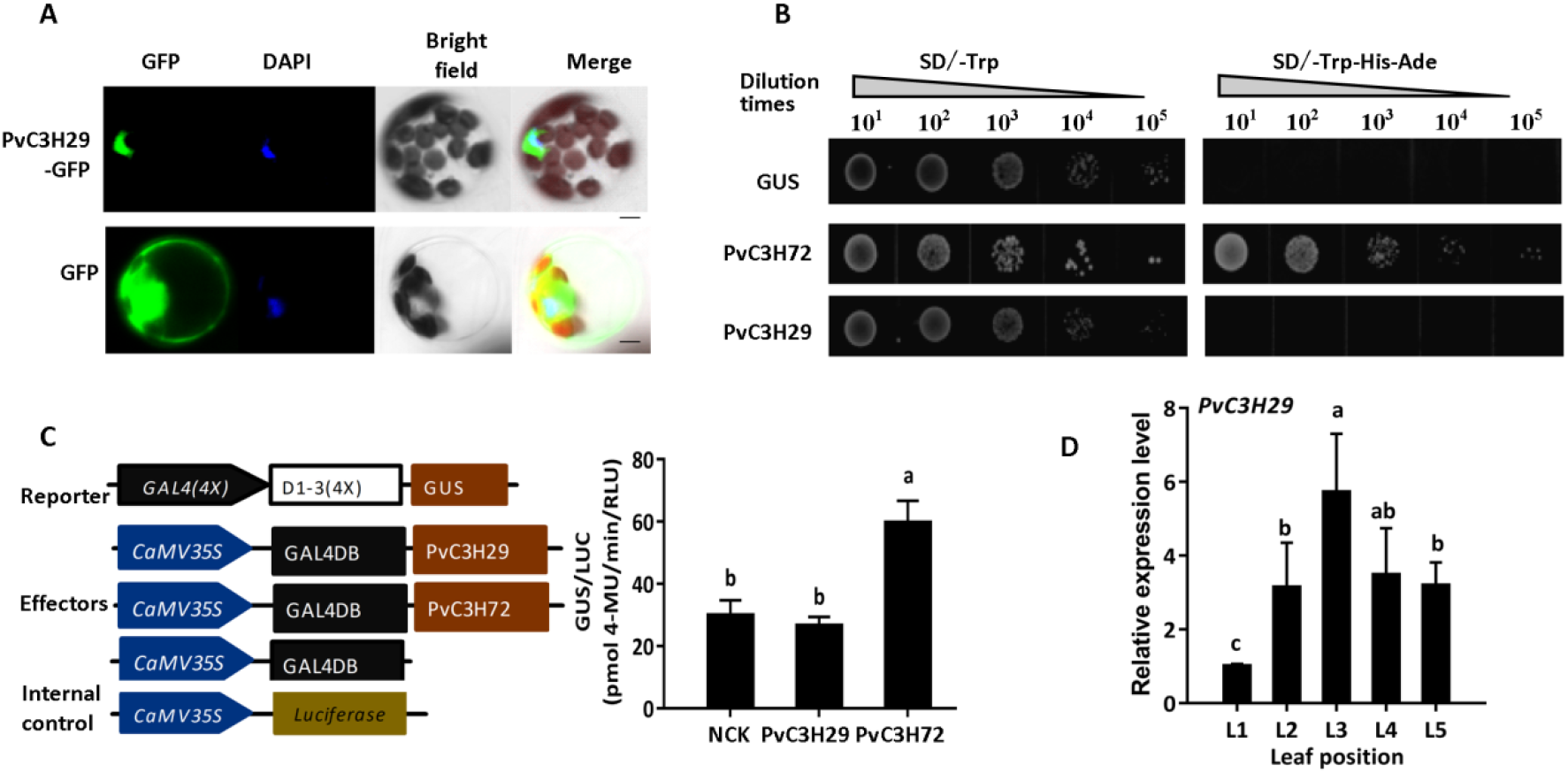
PvC3H29 is a nuclear-localized protein with no transcription activity. A, Subcellular location of PvC3H29-GFP and GFP (control) observed under GFP-fliiorescence, DAPI. and bright field (BF). B, Transcriptional activity assay of PvC3H29 in yeast. PvC3H72, the positive control, was a known transcriptional activator, while the *GUS* gene (*UidA*) as the negative control. C, *In planta* transcriptional activity assay. PvC3H72 was used as the positive control, and the empty vector as the negative control. Letters above bars indicate significant difference at *P* < 0.05. D, Relative expression of *PvC3H29* in leaves from the top at the R3 stage of switchgrass. L1 to L5 were leaves numbered from the top. Bar in (A) represent 5 μm.

### PvC3H29 interacted with PvNAPs in switchgrass

Given that PvC3H29 *per se* had no transcriptional activity but suppressed leaf senescence, there was a possibility that PvC3H29 functioned through protein interaction with senescence-associated transcription factors. To test this hypothesis, we screened its potential interacting proteins with an emphasis on TFs using the Yeast two-hybrid (Y2H) system. After three rounds of library screens, we identified a total of 20 potential interacting proteins (Supplementary Table S1). Among these potential interactors, one pair of paralogous transcription factors were orthologous to the *OsNAP* in rice and were named as *PvNAP1* (Pavir.9KG092600 in ‘P. virgatum v5.1’) and *PvNAP2* (Pavir.9NG600600), respectively. *PvNAP1* and *PvNAP2* are on two homeologous chromosomes (Chr 9a and 9b) of the allotetraploid switchgrass, and their encoded transcriptional factors shared an identical DNA-binding domain (DBD) (Ooka *et al*., 2003) (supplementary Fig. S7), suggesting that they likely targeted on similar set of downstream genes.

Same to that of PvC3H29, subcellular localizations of PvNAP1 and PvNAP2 were in the nucleus (Fig. 4A). The expression of both *PvNAP1* and *PvNAP2* increased along with leaf aging with a similar trend to those of two *CCG*s (*PvSGR* and *PvPAO*) (Fig. 4B). The subcellular localization and expression results met the prerequisite for the physical interaction between PvC3H29 and PvNAP1&2 in switchgrass leaves.

**Fig. 4.**
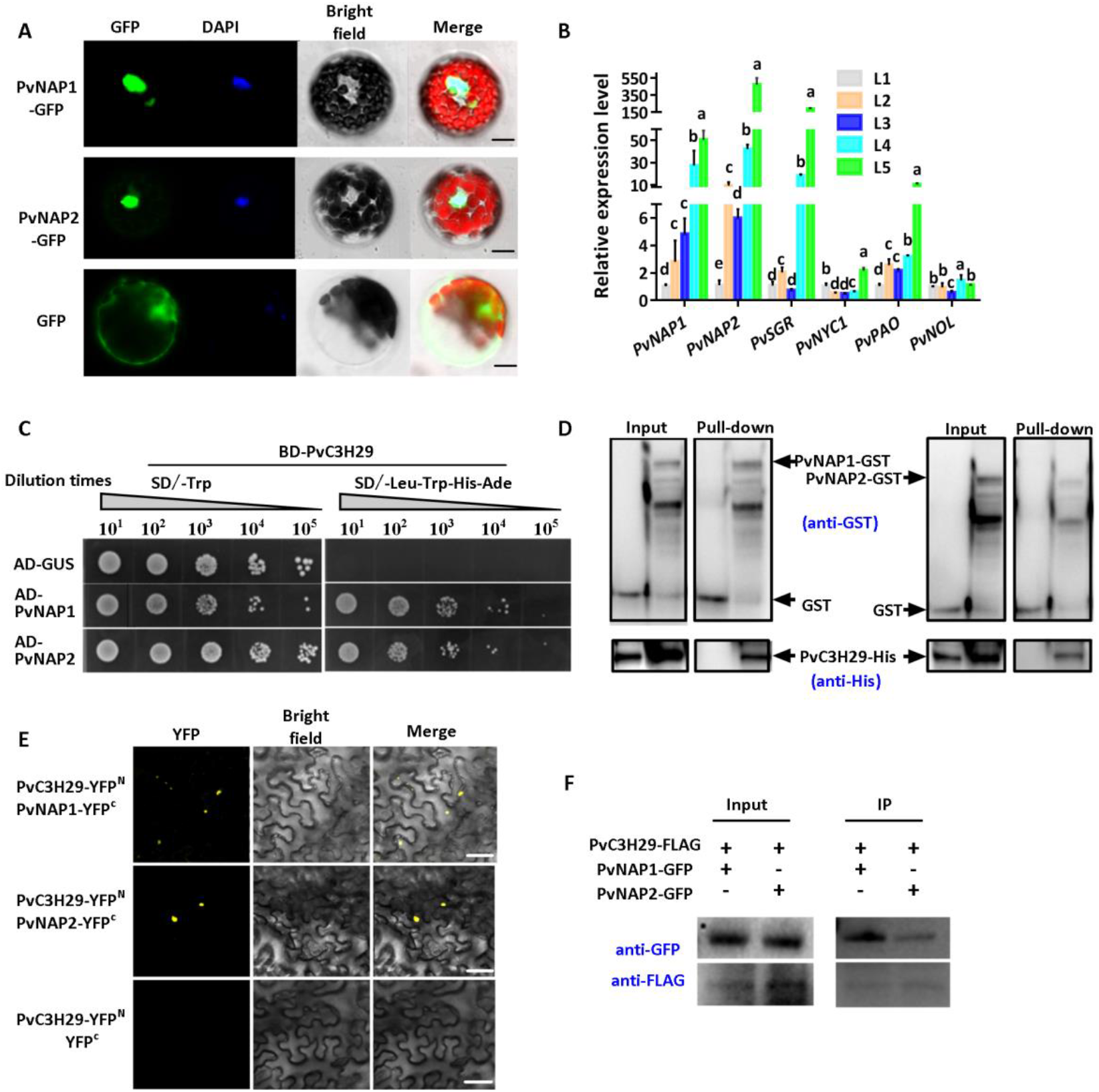
PvC3H29 physically interacts with PvNAP1/2. A, Subcellular location of PvNAP1, PvNAP2, and GFP (control) observed under GFP-fluorescence, DAPI, and bright field (BF). B, Relative expression of *PvNAP1, PvNAP2, PvNYC1, PvNOL, PvSGR*, and *PvPAO* in leaves from the top at the R3 stage of switchgrass. L1 to L5 were leaves numbered from the top. C, Yeast two-hybrid assay of PvC3H29 and PvNAP1/2. D, Pull-down assay between PvC3H29-His and PvNAP1/2-GST. E, BiFC analysis using split YFP with PvC3H29-nYFP and PvNAP1/2-cYFP, and the YFP signal was observed in nucleus. BF stands for bright field. F, Co-IP assay. The protein extract was immunoprecipitated using anti-FLAG. Letters above bars in (B) indicate significant difference at *P*< 0.05. Bars in (A) and (E) represent 5 μm.

To further confirm the physical interaction between PvC3H29 and PvNAPs, we cloned their full CDS and re-tested their interactions by Y2H (Fig. 4C). Secondly, we carried out the pull-down assay and found that the His-tagged PvC3H29 protein could be specifically pulled-down together with the GST-tagged PvNAP1 or PvNAP2 but not with the GST alone (the negative control), confirming their direct interactions *in vitro* (Fig. 4D). Thirdly, we applied the Bimolecular Fluorescence Complementation (BiFC) assay with PvC3H29 infusion with the N-terminal YFP (nYFP) while PvNAPs fused with the C-terminal YFP (cYFP). As shown in figure 4E, when PvC3H29-nYFP was co-expressed with PvNAPs-cYFP in leaves of *Nicotiana benthamiana*, the YFP signal was detected in nuclei, whereas no signal was detected in negative controls. Fourthly, transiently co-expressed PvC3H29-FLAG and GFP-PvNAPs were immunoprecipitated with anti-GFP and A/G magnetic beads and then immunoblotted with anti-FLAG antibody. As shown in figure 4F, PvC3H29-FLAG was co-precipitated together with PvNAPs. Taken together, the Y2H, pull-down, BiFC, and co-Immunoprecipitation (co-IP) analyses proved that PvC3H29 and PvNAPs directly interacted in the nucleus.

### PvC3H29 suppressed PvNAPs-induced precocious leaf senescence

To understand the biological implication of the interaction between PvC3H29 and PvNAPs, we ectopically over-expressed these genes in Arabidopsis. Firstly, consistent with the switchgrass OE29 lines, ectopic over-expressing *PvC3H29* in Arabidopsis also resulted in the ‘staygreen’ trait with significantly delayed leaf senescence and Chl degradation (Supplementary Fig. S8). Using the Y2H system, we found that PvC3H29 also interacted with the Arabidopsis NAP (Supplementary Fig. S8), suggesting that PvC3H29 functioned in a conserved way in switchgrass and Arabidopsis, likely through physical interaction with NAPs. Secondly, we generated dexamethasone-(DEX) inducible *PvNAPs* (OE-*PvNAP1/2*) in Arabidopsis. As shown in figure 5, after DEX induction, the *OE-PvNAP1/2* plants turned precocious leaf senescence with activated expression of *CCG*s (i.e, *NYC1, SGR*, and *PPH*). Thirdly, we co-expressed *PvC3H29* (*35S::PvC3H29*) and *PvNAP1/2* (DEX-inducible) in Arabidopsis and found that *PvC3H29* effectively alleviated the precocious leaf senescence caused by *PvNAP1/2* with suppressed expression levels of *CCG*s (i.e., *NYC1, SGR*, and *PPH*) (Fig. 5). Together, these results suggested that PvC3H29 could alleviate PvNAP1/2*’* effects on precocious leaf senescence.

**Fig. 5.**
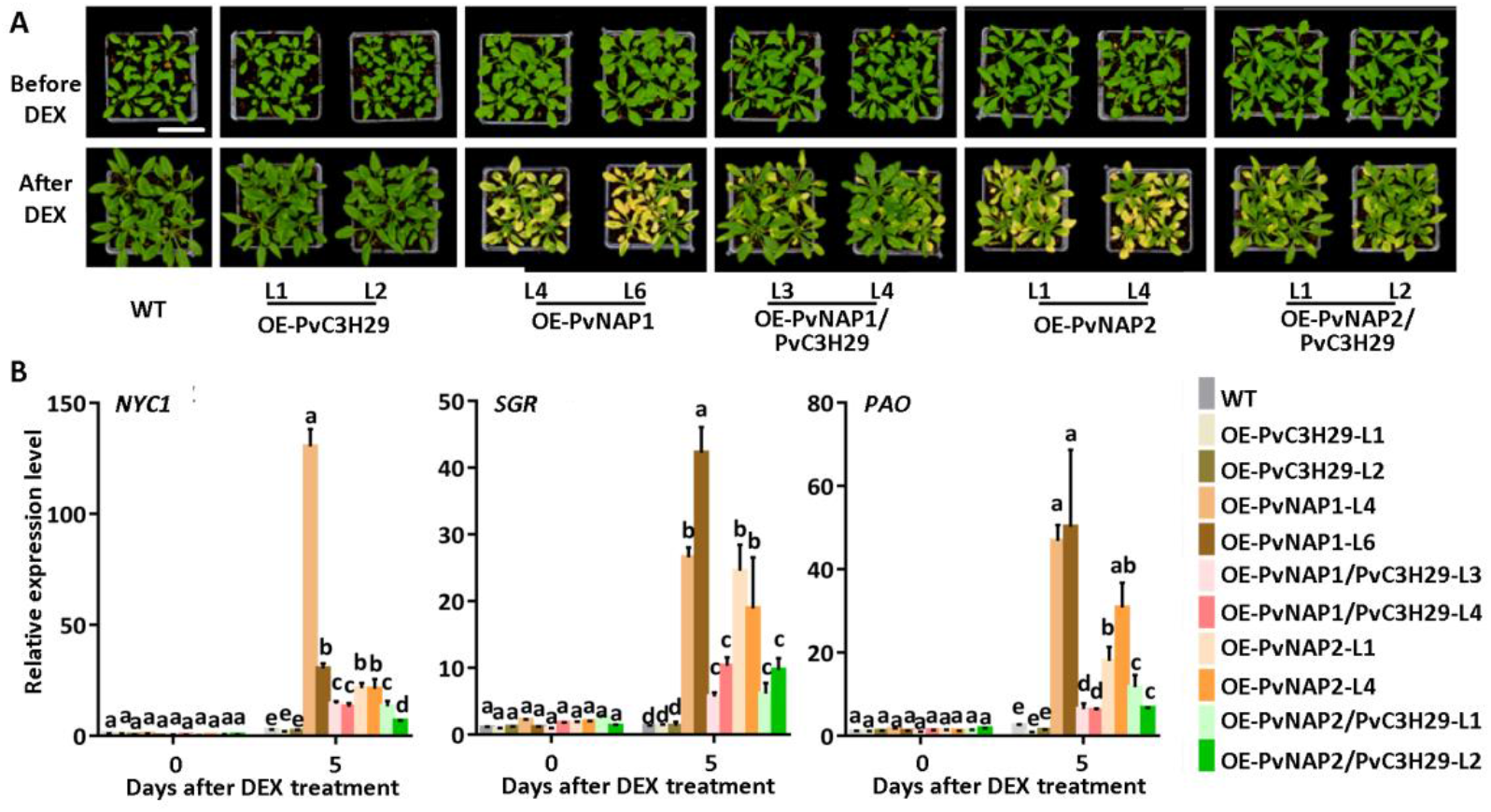
Co-expression of *PvC3H29* suppressed *PvNAP1/2*-induced leaf senescence. **A,** Phenotype of WT and transgenic lines before and after DEX-induced expression of PvNAP1/2. B, Relative expression of *NYC1, SGR*, and *PAO* before and after DEX treatment. Letters above bars in (B) indicate significant difference at *P*< 0.05.

### PvNAPs directly targeted on Chl catabolic genes and transactivated gene expression

As stated above, over-expressing PvC3H29 caused the typical ‘staygreen’ phenotype with repressed *CCG*s, while the contrary was true for PvNAP1/2. Since OsNAP, the rice ortholog to PvNAP1&2, directly targeted on *CCG*s (Liang *et al*., 2014), we further verified whether PvNAPs could directly trans-activate *CCGs*. Since the DNA-binding domains (DBD) of PvNAP1 and PvNAP2 were of identical sequence (supplementary Fig. S7), we used PvNAP1 in the following Yeast one-hybrid (Y1H) and electrophoretic mobility shift assay (EMSA). Firstly, in the Y1H assay, PvNAP1 directly bound promoters of *PvSGR*, *PvPAO*, and *PvNOL* (abbreviated as *pPvSGR*, *pPvPAO*, and *pPvNOL*) (Fig. 6A). Secondly, the EMSA result showed that PvNAP1 caused gel-shift of four probes (E1-E4) of *pPvSGR* (supplementary Fig. S9), and the signal of shifted band decreased with increased concentrations (50×and 200×) of the unlabeled E1 probe (Fig. 6B). Thirdly, *in planta* transactivation analysis by co-expressing the effector (*35S::PvNAP1/2*, or *35S::GFP* for control) and the reporter gene (*LUC*) driven under *pPvSGR,pPvPAO*, or *pPvNOL* showed that both PvNAP1 and PvNAP2 trans-activated these *CCG*s (Fig. 6C). Together, these results showed that PvNAPs directly targeted and transactivated the expression of *PvNOL, PvSGR*, and *PvPAO*.

**Fig. 6.**
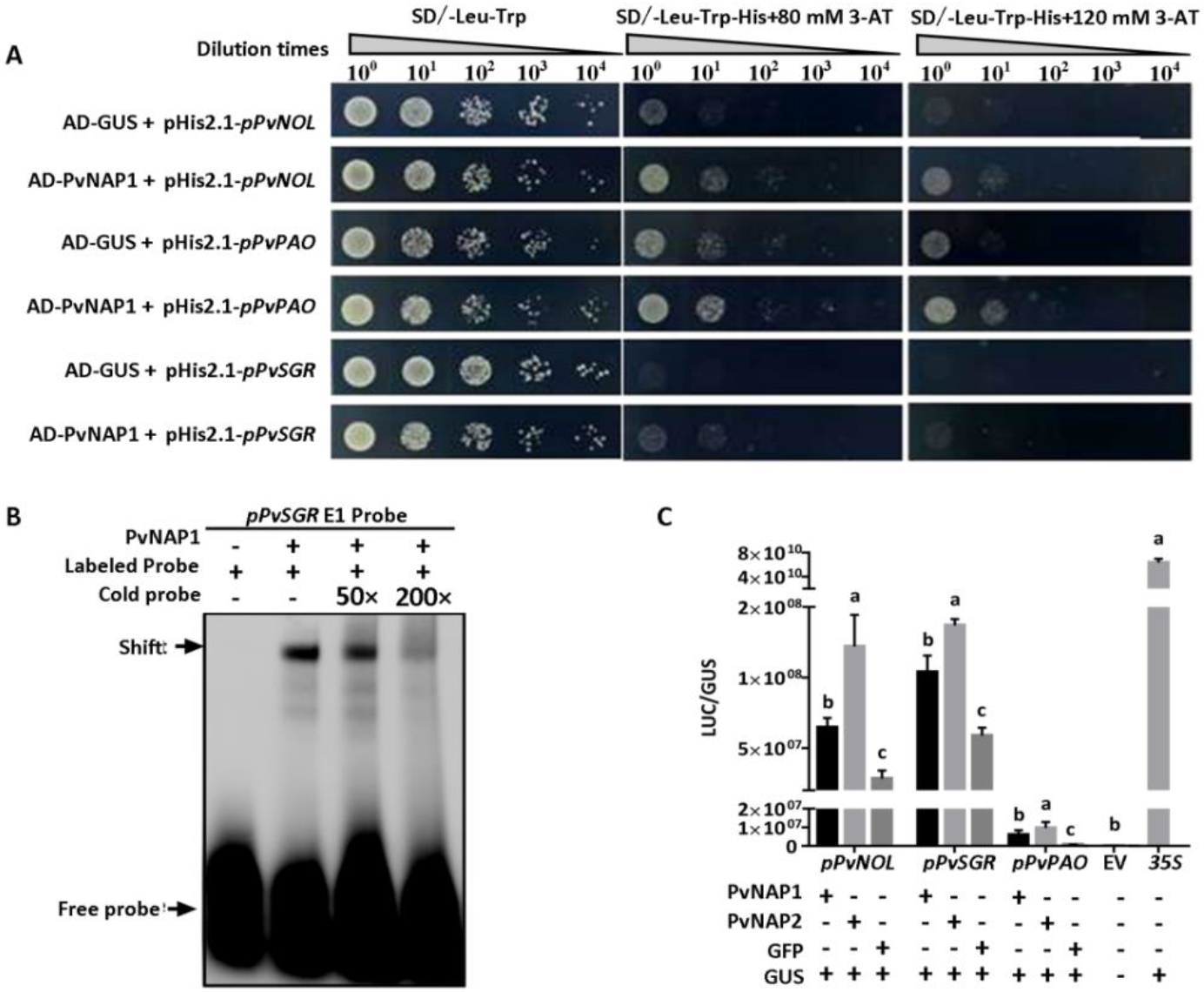
PvNAPs directly targeted and transactivated the expression of *PvNOL, PvSGR*, and *PvPAO*. A, Yeast one-hybrid assay showing that PvNAP1 bounds to promoters of *PvNOL, PvSGR*. and *PvPAO*. GUS was used as the negative control. B, EMSA showing that PvNAP1 directly bound to the promoter of *PvSGR*, C, *In planta* transcriptional activation assay by co-expression of *PvNAP1/2* and *LUC* driven under promoters of *PvNOL, PvSGR*, and *PvPAO*, GFP and empty vector (EV) were used as the negative effector control and the negative *LUC* control, *35S::LUC* as the positive control, and *35S::GUS* as the internal reference. Letters above bars indicate significant difference at *P*< 0.05.

### PvC3H29 inhibits the DNA binding efficiency of PvNAP1

To further understand how PvC3H29 alleviated PvNAPs*’* effects on precocious leaf senescence, we checked whether the interaction with PvC3H29 affected PvNAPs’ DNA binding efficiency using PvNAP1 and *pPvSGR* as the exemplar pair. Using EMSA, we found that PvC3H29 *per se* did not bind the E1 probe, whereas incubation PvNAP1 together with PvC3H29 weakened the signal of shifted E1 probe (Fig. 7A), confirming that PvC3H29 inhibited the DNA binding efficiency of PvNAP1. Furthermore, using the *in planta* transactivation assay, we showed that PvC3H29 significantly suppressed the transactivation of *pPvSGR* by PvNAP1 (Fig. 7B). Taken together, these results proved that PvC3H29 inhibited the DNA-binding of PvNAP1 onto its target genes’ promoters, thereby repressing leaf senescence.

**Fig. 7.**
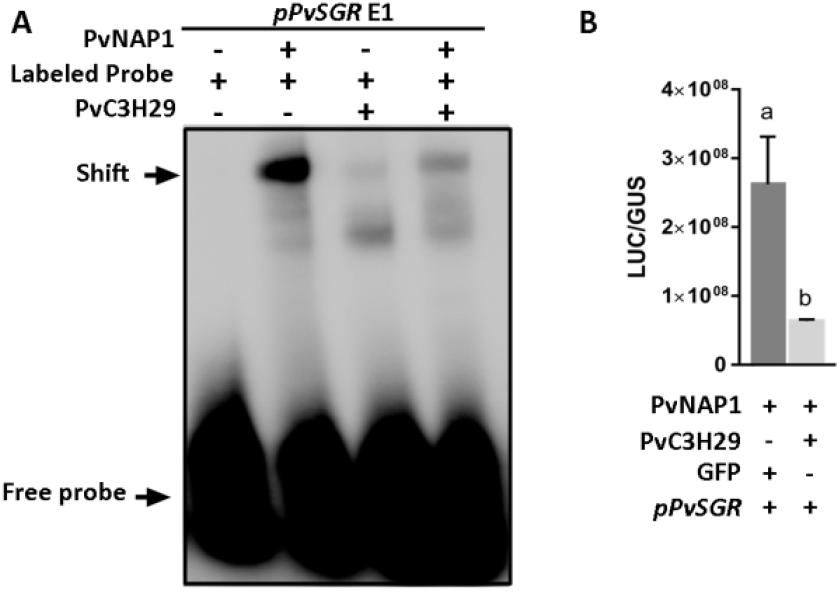
PvC3H29 blocked PvNAP1’s DNA binding efficiency on its target promoter. A, EMSA showing that the addition of PvC3H29 reduced DNA-binding efficiency of PvNAP1 to the *pPvSGR-E1* probe. B, *In planta* transcriptional activation assay showing that co-expression of *PvC3H29* together with *PvNAP1* reduced the expression level of *LUC* driven under *pPvSGR*. GFP was used as the negative effector control, and 35S::*GUS* was used as the internal reference. Letters above bars indicate significant difference at *P*< 0.05.

### Improved biomass yield and feedstock quality by over-expressing *PvC3H29* in switchgrass

Over-expressing *PvC3H29* resulted in functional staygreen phenotype with no apparent growth penalty in the perennial tall grass. When harvested at the R3 stage (Hardin *et al*., 2013), the three switchgrass OE29 lines gained 30%~47% higher biomass yields, 28%~40% higher leaf: stem ratios, 36%~98% more tiller numbers, 31%~66% higher crude protein contents, and 100%~140% higher soluble sugar contents than those of WT (Fig. 8), and their cell wall compositions were similar to that of WT (e.g., NDF, ADF, cellulose, and hemicellulose contents) (Supplementary Fig. S10).

**Fig. 8.**
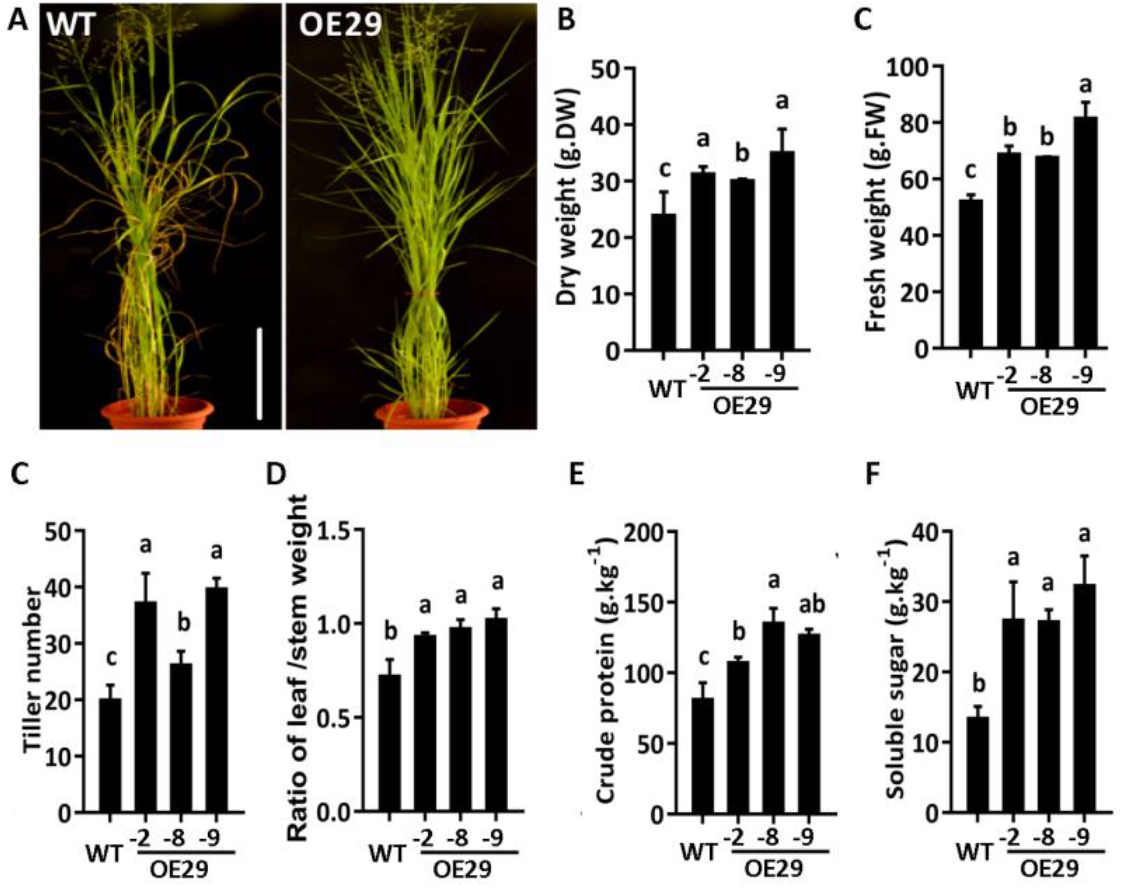
Over-expressing *PvC3H29* improved biomass yield and feedstock quality in switchgrass. A, Phenotype of WT and OE29 at R3 stage. B-F, Dry weight (A), fresh weight (B), tiller number (C), leaf: stem weight ratio (D), crude protein content (E), and soluble sugar content (F) of the above-ground biomass of WT and OE29 lines harvested at R3 stage. Letters above bars indicate significant difference at *P* < 0.05.

## Discussion

Leaf senescence is a complicated process involving extensive reprogramming and gene regulatory network (Quirino *et al*., 2000; Woo *et al*., 2019). If we imagine the progression of leaf senescence as an ‘one way train’, the operator needs both ‘accelerators’ and ‘brakes’ to drive it to the final destination. Previous studies on the transactivation of *CCGs* mainly focused on how senescence-related signals activated *CCGs* through various TFs (Kuai *et al*., 2018). Yet, the inhibitory mechanism to repress *CCGs*’ expression in mature pre-senescing green leaves are not well studied. In this study, we reported a new regulatory module in leaf senescence consisting of a novel CCCH-type zinc finger protein, PvC3H29, and a pair of senescence-promoting transcription factors, PvNAPs. Since either knocking-down *PvC3H29* or ectopic over-expressing *PvNAPs* led to precocious leaf senescence, we proposed that the regulatory module of PvC3H29-PvNAPs were indispensable for the proper rate of leaf aging.

Up to date, there was little report on the interaction between CCCH Znf and NAC TFs and their regulatory link in leaf senescence regulation was missing, although a number of CCCH Znf proteins and NAC TFs are involved in leaf senescence. In this study, we found PvC3H29 had a strong effect on leaf senescence and Chl catabolism. Since PvC3H29 has no transactivation activity *per se,* we then focused on its interacting proteins. The physical interaction between PvC3H29 and the pair of leaf senescence positive TFs (PvNAP1&2) effectively suppressed the TF’s binding efficiency to CCGs’ promoters and thereby repressed Chl degradation and leaf senescence. TZF1, OsDOS, OsTZF1, and PvC3H69 also acted as negative regulators in leaf senescence in Arabidopsis, rice, and switchgrass (Kong *et al*., 2006; Jan *et al*., 2013; Pomeranz *et al*., 2010; Qu *et al*., 2014; *Xie et al., 2021*). The direct interaction between PvC3H29 and PvNAPs might be an exemplar link between the CCCH-type Znfs and NACs to co-regulate Chl degradation and leaf senescence. In the future, it would be interesting to test whether there is a similar regulatory module between senescence-promoting transcription factors and the other CCCH-Znfs.

The paralogous pair of PvNAP1&2 shared the same DNA-binding domain and presumably highly similar promoter-binding preferences, similar expression patterns, and functions in leaf senescence. As stated before, these two paralogs are on homeologous chromosomes (Chr 9a and 9b) of the allotetraploid switchgrass and have exactly the same sequences in the putative DNA-binding domains to keep their conserved functions. The orthologous NAPs seem to have very conserved functions in accelerating leaf senescence not only within the grass family species but across dicot and monocot species. For examples, the rice OsNAP and Arabidopsis NAP both function as positive regulators of leaf senescence but targeting on different set of senescence-related genes that the OsNAP targets on *CCGs* while the Arabidopsis NAP targets on ABA-biosynthesis genes (Yang *et al*., 2014; Liang *et al*., 2014). It is interesting to note that PvC3H29 also interacted with the Arabidopsis NAP that could also explain the ‘staygreen’ phenotype of Arabidopsis OE29 lines. We argue that such a conserved function among orthologous NAPs should reflect their essential roles in the regulation of leaf senescence and the long-term survival of different plant species, considering that the leaf senescence-associated nutrient remobilization to reproduction, rhizome, and other sink organs was one crucial aspect for the success of both annual and perennial plant species. On the other hand, precocious leaf senescence in an ‘uncontrollable manner’ will be detrimental to plants’ fitness as well. Therefore, we propose that there the sophisticated interacting network between the senescence-promoting factors (‘accelerator’, PvNAPs in this case) and senescence-delaying factors (‘brakes’, e.g., PvC3H29).

Over-expressing *PvC3H29* did not result in any obvious drawback in terms of growth penalty or tillering re-growth after cut back for years in our controlled environment. This observation suggested the plasticity of switchgrass genetic manipulation with a promise for the use of the ‘staygreen’ trait. As a bioenergy and animal feedstock crop, biomass yield and feedstock quality of switchgrass are the primary traits for genetic improvement (Anderson *et al*., 1988; McLaughlin and Kszos, 2005). Switchgrass OE29 lines had up to 47% higher biomass yields, 66% higher crude protein contents, and 140% higher soluble sugar contents than those of WT, supporting that delayed leaf senescence is an important aspect to consider for molecular breeding of bioenergy and forage grass. The longer leaf lifespan should have contributed to the longer photosynthetic period and thus accumulation of more biomass, while leaves of longer juvenile stage contributed to the higher soluble sugar and protein contents (Yang *et al*., 2018). Similar findings were also reported in the major grain crop, rice, that natural variation in *SGR* promoters causes rapid Chl degradation and is responsible for the early leaf senescence in indica-type rice (*O. sativa* ssp. *indica)* (Shin*et al*.,2020). Introgression of the *SGR* promoter from the *japonica-type* rice (*O. sativa* ssp. *japonica)* into the *indica*-type rice varieties delayed Chl degradation and leaf senescence and led to 10.6-12.7% increase of grain yield in the near isogenic lines of *indica* rice (Shin *et al*., 2020). Previous studies on *SGR* in alfalfa (Zhou *et al*., 2011) and perennial ryegrass (Xu *et al*., 2019) also showed that the *sgr* mutant or RNAi lines had greener feedstock appearance as well as higher protein contents. OE29 lines showed not only delayed leaf senescence but also improvements in biomass yield and feedstock quality, thus further reiterated the ‘staygreen’ trait in crop improvements.

In sum, PvC3H29 functioned as a repressor in Chl degradation and leaf senescence at least partially through its interaction with PvNAPs to repress their DNA binding efficiency on their downstream genes, such as *CCGs*.

## Materials and Methods

### Gene cloning and vector construction

The CDS of *PvC3H29*, *PvNAP1* and *PvNAP2* was amplified from cDNA of a selected line ‘HR8’ of the tetraploid switchgrass ecotype ‘Alamo’ (Xu *et al*., 2011a), cloned into the Gateway entry vector pENTR/D (Invitrogen Life Technologies, Carlsbad, CA, USA), and then subcloned into destination vectors, such as p2GWF7.0 for subcellular localization assay (Karimi *et al*., 2002), pGBKT7 and pGADT7 (Invitrogen) for yeast two hybrid and transactivity assays, pVT1629 (Xu *et al*., 2011a), pEarlygate103, and pTA7001 for genetic transformation of switchgrass or *Arabidopsis*. A unique fragment of PvC3H29 were selected for RNAi (supplementary Table S2), and subcloned into the RNAi vector, pGM-kannibal, to form the hairpin structure, and then the hairpin structure was subcloned to the destination vector, pVT1629. The primers used for cloning and vector construction are shown in supplementary Table S3.

### Switchgrass and *Arabidopsis* genetic transformation

Switchgrass genetic transformation was carried out following the protocol described in Xu *et al*. (2011b). In brief, embryogenic calluses of the switchgrass line ‘HR8’ was used for transformation, selected on 50 mg L^-1^ hygromycin (Sigma). Plants generated from different calluses were considered as independent transgenic events. GUS staining and PCR for the presence of the T-DNA fragment of transgenic lines were the same as described in Xu *et al*. (2011a).

*Arabidopsis* transformations were performed using the floral dip method (Zhang *et al*., 2006). *PvNAP1/2* were subcloned into the dexemethesone (DEX)-inducible vector pTA7001 (Aoyama and Chua,1997). PvC3H29 were subcloned into pEearlygate101 (Earley *et al*., 2006). Then, the resultant vectors in *A. tumefaciens* strain ‘AGL1’ were individually or co-transformed to *Arabidopsis*.

### Plant growth conditions and dark treatment

Switchgrass plants were grown in a greenhouse with temperatures set at 28/22°C (day /night) and a 14-h light/10-h darkness. The plants were watered twice a week. In order to induce leaf senescence, the middle section of fully-expand leaves from three-month-old plants were cut into 3 cm segments and placed in dampened paper towels in a dark room with air temperature controlled at 28°C.

*Arabidopsis* ecotype ‘Columbia-0’ was used in this study. Seeds were surface sterilized and grown on ½ Murashige and Skoog (MS) medium with 3% phytogel, then stratified for 3 days at 4°C to synchronize germination. Seedlings were transplanted to soil and grown in a growth chamber at 25/20°C (day /night), light intensity of 120 μmol·m^-2^·s^-1^ under a 16-h light/8-h dark photoperiod.

### Feedstock quality analysis

Four-month-old plants were used for phenotypic and physiological analysis of natural senescence. The whole aboveground WT and over-expression plants were collected for Fresh weight and then dried in a 70°C oven for two weeks for Dry weight and feedstock Quality analysis. Total soluble sugar was determined by the phenol sulfuric acid reagent method according to Dubois *et al*. (1956). Cellulose, hemicellulose, ADF and NDF were determined in reference to Goering and Van Soest (1970). The analysis of crude protein content was according to a protocol described by Janicki and Stallings (1987). Leaf /steam was calculated by the radio of leaf dry weight and steam dry weight.

### Chl content and Fv/Fm measurements

For Chl content measurement, leaves were soaked in dimethylsulfoxide (DMSO) for 48h in dark, and the extract was measured at 663nm and 645nm (Barnes *et al*., 1992) using a spectrophotometer (Spectronic Instruments, Rochester, NY, USA).

For Fv/Fm measurement, leaves were adapted to dark condition for 30 min prior to the measurement, then the Fv/Fm values were measured using a fluorescence meter (Dynamax, Houston, TX, USA).

### Subcellular localization

*PvC3H29*, *PvNAP1* and *PvNAP2* were subcloned into a modified gateway compatible P2GWF7.0 vector (Karimi *et al*., 2002) to generate GFP fusion genes driven under *CaMV* 35S promoter. The PvC3H29-GFP, PvNAP1-GFP and PvNAP2-GFP fusion genes were transiently expressed in *Arabidopsis* protoplasts through polyethylene glycol (PEG)-mediated protoplast transformation (Yoo *et al*., 2007). DAPI was used to stain the nucleus, and the GFP signal was detected under a Zeiss LSM 780 laser scanning confocal microscope (Carl Zeiss SAS, Jena, Germany).

### Transcriptional activity assay

For the yeast-based auto-transactivation assay, the CDS of *PvC3H29, UidA*(GUS-encoding gene, negative control), *PvC3H72* (positive control) was subcloned into pGBKT7 to fuse the gene with the DNA binding domain of *GAL4*. The generated vectors were transformed into the yeast strain Y2HGold (Clonetech Laboratories, Palo Alto, CA, USA), separately. The PvC3H72 was a known transcription activator(Xie *et al*., 2019). The transformed positive clones were selected on SD/-Trp, which were then grown on plates containing SD/-Trp-Ade-His and SD/-Trp-Ade-His + 25 mM 3-AT for auto-transactivation assay.

*In planta* transcriptional activity assay was the same as described previously (Xie *et al*., 2019). *PvC3H29* were subcloned into the effector vector infusion with *GAL4DB*, while *PvC3H72* and *UidA* genes were used as positive and negative controls, respectively.

### Real time quantitative reverse transcription PCR (RT-qPCR)

To analyze the relative expression levels of *PvC3H29, PvNAP1&2, PvNYC1, PvNOL*, and *PvPAO* in leaves at different developmental stages, we collected the leaves of different orders (i.e., the flag leaf as the 1^st^ and followed by the 2^nd^ to the 5^th^ leaves from the top) in switchgrass at the “R3” stage featured with fully emerged spikelets and an emerged peduncle (Hardin *et al*., 2013). For the analysis of the staygreen trait among WT, OE29, and RNAi lines, we sampled detached leaves after different days of dark treatment for the RT-qPCR analysis. Total RNA extraction, PCR reaction and data analysis were the same as described before (Xie *et al*., 2019). Primers used in this study were listed in supplementary table S3.

### Transcriptome comparison among WT, OE29, and RNAi plants

RNA-Seq was carried out using the paired-end technology of the Illumina HiSeq™ 2000 platform, and the data can be achieved at NCBI (SRA accession: PRJNA780344). Transcriptomic data analysis were performed by Gene Denovo Co. (Guangzhou, China) following the same procedures as reported before (Xu et al., 2019). Detached leaves at the same developmental stage were placed in dampened paper towels in a dark room with air temperature controlled at 28°C for seven days. Leaves from one plant were regarded as one biological sample, and three independent WT, OE29, and RNAi lines were used. The RPKM values (Number of uniquely mapped reads per kilobase of exon region per million mappable reads) were used for identifying differentially expressed genes (DEGs) based on the threshold of FDR=0.001 and an absolute value of log_2_ (RPKM of WT / RPKM of OE29 or RNAi) ≥ 1.

### Yeast two-hybrid

The CDS of *PvC3H29* was subcloned into the vector PGBKT7, and the resultant vector was transformed into the yeast strain ‘Y2H gold’. A cDNA library constructed using switchgrass leaves’ cDNA were screened following the Matchmaker 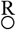 Gold Y2H system user manual (Clontech). The library screening was performed on SD/-Trp-Leu-Ade-His medium. To verify the putative positive clones (preys) from the library screening, we cloned the full length CDS of the preys first, subcloned them into pGADT7, and then tested their interactions with PvC3H29 one by one in the Y2H system again.

### Bimolecular fluorescence complementation assay (BiFC)

The BiFC analysis, PvNAP1/2 and PvC3H29 were subcloned into pFGC-YC155 and pFGC-Y173 vectors (Lai *et al*., 2011). The resultant vectors, harboring the target genes in fusion with C-terminal or N-terminal of YFP, respectively, were transformed into *A. tumefaciens* strain ‘EHA105’, and then transiently co-infiltrated into the leaves of *N. benthamiana*. The images were captured using a confocal laser-scanning microscope (Carl Zeiss SAS, Jena, Germany).

### Yeast one-hybrid

The recombined pGADT7 vectors (pGADT7-PvNAP1, pGADT7-PvNAP2 and pGADT7) and pHIS2.1 vectors (pHIS2.1-*pPvNOL*, *-pPvPAO*, and *-pPvSGR*) were co-transformed into the yeast strain *Y187*. After three days growth, five clones for each plate were picked, grown in liquid medium for three days at 30°C and then dotted on solid media of SD/-Trp-Leu and SD/-Trp-Leu-His with 50, 80 and 120 mM 3-AT, respectively.

### Transient transactivation assay in *N. benthamiana*

Promoters of four *CCGs* (*pPvNOL*, *pPvPAO*, and *pPvSGR*), were subcloned into the pCambia1381Z reporter vector. PvNAP1/2 and PvC3H29 were cloned intopEarleyGate103as the effector vector with pEarleyGate103-*eGFP*as the negative control. pCambia1302-*GUS-polyA* was used as the internal control with *GUS* driven by *CaMV 35S* promoter. For the activating effect of PvNAP1/2 on *CCGs*, the reporter, effecter, and internal control were mixed at the ratio of 1:1:1. While for the test on repressing effect of PvC3H29 on PvNAP1/2, we used the ratio of 3:3:2:1 for each effector, reporter and internal control, respectively. Injected leaves were sampled after two days and ground with liquid nitrogen. The soluble proteins were extracted with a extraction buffer (Yeasen Biotech, Shanghai, China) for LUC and GUS activity assays using kits according to the providers’ protocols (Coolaber, Beijing, China).

### Pull-down assay

For protein expression and purification, the GST-PvNAP1/2 fusion proteins were generated using the pGEX4T-1vectorand the recombinantHis-PvC3H29 fusion protein was generated using the pCold™ TF vector and expressed in the *E. coli* strain *Bl21* after isopropylb-D-1-thiogalactopyranoside (IPTG)induction at 16°C. Recombinant proteins were purified using GST Resin and Ni Resin, respectively (Transgene, Beijing, China).For Pull-down assay, GST-PvNAP1/2 or GST proteins(100μg) was incubated with glutathione-Sepharose4Bresin(Transgene)at 4°Cfor one hour to make the beads fully linked with the GST-PvNAP1/2 or GST proteins and then washed three times with 500 μl Lysing buffer. Then, His-PvC3H29 protein at the same volume was incubated with GST-PvNAP1/2 or GST linked beads for two hours. The conjugated constructs were centrifuged and washed three times with lysing buffer to prepare for the immunoblot analysis. Finally, the protein extract before and after immunoprecipitation were analyzed using both anti-GST (Transgene, Beijing, China) and anti-His antibodies (Transgene).

### Co-IP

*PvC3H29* was subcloned into pEarlygate202 vector to generate PvC3H29-FLAG fusion protein, while *PvNAP1/2* were into pEarlygate103 to generate PvNAP1/2-GFP fusion proteins. These vectors were transiently co-expressed in *N. benthamiana* leaves. After 48 h, leaves were homogenized in extraction buffer containing 50mM Tris-HCl (pH 7.5), 150 mM NaCl, 1 mM EDTA, 0.1% (w/v) Triton X-100, and 1×complete protease inhibitor cocktail (Roche). Then, the protein extract was immunoprecipitated using an anti-FLAG antibody (Thermo Scientific, USA), and the co-immunoprecipitated proteins were detected using an anti-GFP antibody (Thermo scientific).

### DEX treatment

For the inducible expression of PvNAP1/2 in *Arabidopsis*, 30 μM DEX was used to spray the whole *Arabidopsis* plant. Leaf samples was taken prior to or 5 days after the treatment for measurements of Chl content, Fv/Fm and for RNA extraction.

### EMSA

pCold™ TF-PvNAP1 was constructed to generate HIS-PvNAP1 fusion gene. The recombinant proteins were expressed and purified using the same protocol as described in the Pull-down assay. Four candidate probes were firstly screened and one of the highly shifted probes was used for the next competitive analysis by following the protocol in the Light Shift Chemiluminescent EMSA Kit (Thermo Scientific). Unlabeled competitors were added in a 50- or 100-fold molar excess. The detect assay was the same as previously reported (Yu *et al*., 2022). As for the repressing activity of PvC3H29 in PvNAP1 binding to *pPvSGR*, three-fold content of His-PvC3H29 was pre-incubated with His-PvNAP1.

### Statistical analysis

Data collected in this study were analyzed using SAS v9.2 (SAS Institute, Cary, NC, USA), and were represented as mean ± standard error (SE). Fisher’s protected LSD was used to discriminate significant difference among sets of data at the probability of 0.05.

## Acknowledgements

We thank Dr. Binyu Zhao from Virginia Polytechnic Institute and State University for providence of several vectors used in the study. This study was funded by National Natural Science Foundation of China (NSFC) projects (31971757 and 31772659).

## Author contributions

BX and GY designed the experiments; ZX and GY conducted the experiment; HW and SL assisted with some of the experiments. ZX analyzed data; BX and ZX wrote the manuscript.

## Supporting Information

**Supplementary Fig. S1 GUS staining and RT-qPCR of *PvC3H29* over-expression and RNAi transgenic lines. (A)** GUS staining of WT, OE29, and RNAi transgenic. **(B)** Relative expression of *PvC3H29* lines in leaves of WT, OE29, and RNAi lines by RT-qPCR. Letters above bars indicate significant difference at *P* < 0.05.

**Supplementary Fig. S2 Over-expressing *PvC3H29* delayed natural leaf senescence in switchgrass. (A)** Leaves of WT and OE29 at R3 stage showing different degrees of senescence. Leaves are numbered from the top. **(B-C)** Chl contents and Fv/Fm values of the leaves. Letters above bars indicate significant difference at *P* < 0.05.

**Supplementary Fig. S3 Number of differentially expressed genes (DEGs) in OE 29 and RNAi transgenic lines.** Detached leaves **treated** in dark for seven days were sampled for the RNA-seq analysis.

**Supplementary Fig. S4 Gene Ontology (GO) analysis of DEGs. (A)** GO terms of WT vs OE29; **(B)** GO terms of WT vs RNAi. Detached leaves treated in dark for seven days were sampled for the RNA-seq analysis.

**Supplementary Fig. S5 Kyoto Encyclopedia of Genes and Genomes (KEGG) analysis of DEGs. (A)** Enriched KEGG pathways in WT vs OE29; **(B)** Enriched KEGG pathways in WT vs RNAi. Detached leaves treated in dark for seven days were sampled for the RNA-seq analysis.

**Supplementary Fig. S6 Comparison between WT, OE29, and RNAi line for Chl catabolic pathway genes.** Green arrows indicate down-regulated expression, while red arrows indicate up-regulated expression in OE29 or RNAi lines. Detached leaves treated in dark for seven days were sampled for the RNA-seq analysis.

**Supplementary Fig. S7 Ectopic over-expressing *PvC3H29* leads to stay-green in Arabidopsis. (A)** Relative expression of *PvC3H29 in* Arabidopsis transgenic lines. Not detectable is abbreviated as n.d. **(B-C)** OE29 Arabidopsis lines showed stay-green after 10 <u>d</u>ays <u>a</u>fter <u>d</u>ark treatment (DAD), and the Chl contents in leaves of WT and transgenic lines during the dark treatment. **(D)** PvC3H29 interacted with the Arabidopsis NAP in the Y2H assay.

**Supplementary Fig. S8 Electrophoretic mobility shift assay (EMSA) of PvNAP1’s binding to four probes in the promoter of *PvSGR*.**

**Supplementary Fig. S9 Neutral and acid detergent fiber (NDF and ADF), cellulose and hemicellulose contents of aboveground biomass in WT and OE29 lines of switchgrass harvested at R3 stage.**

**Supplementary Table S1.** Potential interacting proteins of PvC3H29 screened through Yeast-two-hybrid (Y2H).

**Supplementary Table S2.** Fragment of PvC3H29 were selected for RNAi.

**Supplementary Table S3.** Primers used in this study.

## References

Anderson B, Ward JK, Vogel KP, Ward MG, Gorz HJ, Haskins FA (1988) Forage quality and performance of yearlings grazing switchgrass strains selected for differing digestibility. JAS 66: 2239–2244

Aoyama T, Chua NH (1997) A glucocorticoid-mediated transcriptional induction system in transgenic plants. Plant J 11: 605–612

Barnes JD, Balaguer L, Manrique E, Elvira S, Davison AW (1992) A reappraisal of the use of DMSO for the extraction and determination of chlorophylls a and b in lichens and higher plants. Environ Exp Bot 32: 85–100

Chen Y, Qiu K, Kuai B, Ding Y (2011). Identification of an NAP-like transcription factor BeNAC1 regulating leaf senescence in bamboo (*Bambusa emeiensis* ‘Viridiflavus’). Physiol Plant 142: 361–371

De Lucas M, Daviere JM, Rodríguez-Falcón M, Pontin M, Iglesias-Pedraz JM, Lorrain S, Fankhauser C, Blázquez MA, Titarenko E, Prat, S (2008) A molecular framework for light and gibberellin control of cell elongation. Nature 451: 480–484

Dubois M, Gilles KA, Hamilton JK, Rebers PT, Smith F (1956) Colorimetric method for determination of sugars and related substances. Anal Chem 28: 350–356

Earley KW, Haag JR, Pontes O, Opper K, Juehne T, Song K, Pikaard CS (2006) Gateway-compatible vectors for plant functional genomics and proteomics. Plant J 45: 616–629

Fan K, Bibi N, Gan S, Li F, Yuan S, Ni M, Wang M, Shen H, Wang X (2015) A novel NAP member GhNAP is involved in leaf senescence in *Gossypium hirsutum*. J Exp Bot. 66: 4669–4682

Goering HK, Van Soest PJ (1970) Forage fiber analyses (apparatus, reagents, procedures, and some applications), US ARS pp: 387–598

Hardin CF, Fu C, Hisano H, Xiao X, Shen H, Stewart CN, Parrott W, Dixon RA, Wang ZY (2013) Standardization of switchgrass sample collection for cell wall and biomass trait analysis. BioEnerg Res 6:755–762

Jan A, Maruyama K, Todaka D, Kidokoro S, Abo M, Yoshimura E, Shinozaki K, Nakashima K, Yamaguchi-Shinozaki K (2013) OsTZF1, a CCCH-tandem zinc finger protein, confers delayed senescence and stress tolerance in rice by regulating stress-related genes. Plant Physiol 161: 1202–1216

Janicki F, Stallings C (1987) Nitrogen fractions of alfalfa silage from oxygen-limiting and conventional upright silos. J Dairy Sci 70: 116–122

Karimi M, Inzé D, Depicker, A (2002) GATEWAY™ vectors for Agrobacterium-mediated plant transformation. Trends Plant Sci 7: 93–195

Kong Z, Li M, Yang W, Xu W, Xue, Y (2006) A novel nuclear-localized CCCH-type zinc finger protein, OsDOS, is involved in delaying leaf senescence in rice. Plant Physiol 141: 1376–1388

Kuai B, Chen J, Hörtensteiner S (2018) The biochemistry and molecular biology of chlorophyll breakdown. J Exp Bot 69: 751–767

Lai Z, Li Y, Wang F, Cheng Y, Fan B, Yu JQ, Chen Z (2011) *Arabidopsis* sigma factor binding proteins are activators of the WRKY33 transcription factor in plant defense. Plant Cell 23: 3824–3841

Lei W, Li Y, Yao X, Qiao K, Wei L, Liu B, Zhang D, Lin H (2020) NAP is involved in GA-mediated chlorophyll degradation and leaf senescence by interacting with DELLAs in *Arabidopsis*. Plant Cell Rep 39: 75–87

Liang C, Wang Y, Zhu Y, Tang J, Hu B, Liu L, Ou S, Wu H, Sun X, Chu J et al (2014) OsNAP connects abscisic acid and leaf senescence by fine-tuning abscisic acid biosynthesis and directly targeting senescence-associated genes in rice. Proc Natl Acad Sci USA 111:10013–10018

McLaughlin SB, Kszos LA (2005) Development of switchgrass *(Panicum virgatum)* as a bioenergy feedstock in the United States. Biomass Bioenergy 28: 515–535

Ooka H, Satoh K, Doi K, Nagata T, Otomo Y, Murakami K, Matsubara K, Osato N, Kawai J, Carninci P et al (2003) Comprehensive analysis of NAC family genes in *Oryza sativa* and *Arabidopsis thaliana*. DNA Res 10:239–247

Pomeranz MC, Hah C, Lin PC, Kang SG, Finer JJ, Blackshear PJ, Jang JC (2010) The *Arabidopsis* tandem zinc finger protein AtTZF1 traffics between the nucleus and cytoplasmic foci and binds both DNA and RNA. Plant Physiol 152: 151–165

Qu J, Kang SG, Wang W, Musier-Forsyth K, Jang JC (2014) The *Arabidopsis* thaliana tandem zinc finger 1 (AtTZF1) protein in RNA binding and decay. Plant J 78: 452–467

Quirino BF, Noh YS, Himelblau E, Amasino RM (2000) Molecular aspects of leaf senescence. Trends Plant Sci 5:278–282

Sakuraba Y, Jeong J, Kang MY, Kim J, Paek NC, Choi G (2014) Phytochrome-interacting transcription factors PIF4 and PIF5 induce leaf senescence in *Arabidopsis*. Nat Commun 5: 4636

Shin D, Lee S, Kim TH, Lee JH, Park J, Lee J, Lee JY, Cho LH, Choi JY, Lee W et al (2020) Natural variations at the Stay-Green gene promoter control lifespan and yield in rice cultivars. Nat Commun 11:1–11

Thomas H., Howarth C.J. (2000) Five ways to stay green. J Exp Bot 51:329–337.

Woo HR, Kim HJ, Lim PO, Nam HG (2019) Leaf senescence: systems and dynamics aspects. Annu Rev Plant Biol 70:347–376

Xie Z, Lin W, Yu G, Cheng Q, Xu B, Huang B (2019) Improved cold tolerance in switchgrass by a novel CCCH-type zinc finger transcription factor gene, *PvC3H72*, associated with ICE1-CBF-COR regulon and ABA-responsive genes. Biotechnol Biofuels 12:1–11

Xie Z, Yu G, Lei S, Zhang C, Xu B, Huang B (2021) CCCH protein-PvCCCH69 acted as a repressor for leaf senescence through suppressing ABA-signaling pathway. Hortic Res 8: 1–14

Xu B, Yu G, Li H, Xie Z, Wen W, Zhang J, Huang B (2019) Knockdown of *STAYGREEN* in perennial ryegrass (*Lolium perenne* L.) leads to transcriptomic alterations related to suppressed leaf senescence and improved forage quality. Plant Cell Physiol 60: 202–212

Xu B, Escamilla-Treviño LL, Sathitsuksanoh N, Shen Z, Shen H, Zhang PY, Dixon RA, Zhao B (2011a) Silencing of *4-coumarate: coenzyme A ligase* in switchgrass leads to reduced lignin content and improved fermentable sugar yields for biofuel production. New Phytol 192: 611–625

Xu B, Huang L, Shen Z, Welbaum GE, Zhang X, Zhao B (2011b) Selection and characterization of a new switchgrass (*Panicum virgatum* L.) line with high somatic embryonic capacity for genetic transformation. Sci. Hortic 129: 854–861

Yang J, Worley E, Udvardi M (2014) A NAP-AAO3 regulatory module promotes chlorophyll degradation via ABA biosynthesis in *Arabidopsis* leaves. Plant Cell 26: 4862–4874

Yang J. Udvardi M (2018) Senescence and nitrogen use efficiency in perennial grasses for forage and biofuel production. J Exp Bot 69: 855–865

Yoo SD, Cho YH, Sheen J (2007) *Arabidopsis* mesophyll protoplasts: a versatile cell system for transient gene expression analysis. Nat Protoc 2: 1565

Yu G, Xie Z, Zhang J, Lei S, Lin W, Xu B, Huang B (2021) NOL-mediated functional stay-green traits in perennial ryegrass *(Lolium perenne* L.) involving multifaceted molecular factors and metabolic pathways regulating leaf senescence. Plant J 106: 1219–1232

Yu G, Xie Z, Lei S, Li H, Xu B, Huang B (2022) LpNAL delays leaf senescence by transcriptional repression of two chlorophyll catabolic genes, *LpSGR* and *LpNYC1*, in perennial ryegrass. Plant Physiol https://doi.org/10.1093/plphys/kiac070

Zhang X, Henriques R, Lin SS, Niu QW, Chua NH (2006) Agrobacterium-mediated transformation of *Arabidopsis thaliana* using the floral dip method. Nat Protoc 1: 641

Zhou C, Han L, Pislariu C, Nakashima J, Fu C, Jiang Q, Quan L, Blancaflor EB, Tang Y, Bouton JH et al (2011) From model to crop: functional analysis of a *STAY-GREEN* gene in the model legume *Medicago truncatula* and effective use of the gene for Alfalfa improvement. Plant Physiol 157:1483–1496

Zhou T, Yang X, Wang L, Xu J, Zhang X (2014) GhTZF1 regulates drought stress responses and delays leaf senescence by inhibiting reactive oxygen species accumulation in transgenic *Arabidopsis*. Plant Mol Biol 85: 163–177

